# Functional Specialization for Language Processing in Inferior Frontal Regions During Early Childhood: Evidence from functional near-infrared spectroscopy individual functional channels of interest approach

**DOI:** 10.1101/2025.02.09.637301

**Authors:** Haolun Luo, Tao Yu, Qun Li, Li Sheng

## Abstract

**Significance:** Early language acquisition represents a fundamental achievement in cognitive development, yet the neural mechanisms underlying this process remain debated, particularly whether specialized language circuits exist from early life or emerge gradually through development.

**Aim:** To investigate functional specialization for language processing in bilateral inferior frontal regions during early childhood, examining whether language-selective regions are distinct from domain-general cognitive networks in toddlers aged 2-4 years.

**Approach:** Using functional near-infrared spectroscopy (fNIRS) with an innovative functional Channel of Interest (fCOI) approach, we conducted two experiments involving adults (N=20) and toddlers (N=22, ages 2-4 years) who completed language processing and cognitive control tasks.

**Results:** We demonstrated early functional specialization in the language-selective region of left inferior frontal gyrus, which showed selective responses to linguistic content while remaining insensitive to cognitive demand manipulations in both age groups. However, language selectivity in the homologous right hemisphere region was present only in adults. The multiple demand regions showed complementary patterns, with right-hemispheric selectivity for cognitive control emerging early.

**Conclusions:** These findings provide evidence for early neural specialization of language processing in the left hemisphere, while revealing ongoing development in right hemispheric organization. Our results support models of early language-specific neural circuits rather than gradual differentiation from domain-general mechanisms, while highlighting the protracted development of language organization.

## Introduction

Language acquisition in early childhood represents a remarkable achievement of human cognitive development, occurring naturally through exposure without formal training^1^. A fundamental question in developmental cognitive neuroscience is how the brain supports early language acquisition. One of the most debated aspects of language development is whether the brain contains specialized circuits for language processing from early in life, or whether these circuits emerge gradually over development. Specifically, there are two competing hypotheses about the neural mechanisms underlying language processing in young children. One possibility is that language processing relies on domain-specific neural mechanisms from early on, suggesting an early neural specialization for language that is distinct from other cognitive functions^2,3^. Alternatively, language processing might initially emerge through domain-general cognitive processes, such as working memory and executive function, with specialized language circuits developing gradually through experience^4,5^.

Studies in adults have identified two distinct neural networks potentially relevant for understanding language development: the ‘core language network’ and the ‘multiple demand’ (MD) network. The language network, which includes regions primarily in left frontal and temporal areas, shows consistent engagement during language processing with limited activation during non-linguistic tasks^6–8^. In contrast, the MD network, comprising bilateral frontal, parietal, cingulate, and insular regions, responds to increased cognitive demands across various cognitive tasks^9,10^. The relationship between these networks and their respective roles in language processing have been debated. Some researchers argue that language processing relies substantially on domain-general cognitive mechanisms implemented in the MD network^11,12^, while others propose more distinct functional roles^13^. Recent adult neuroimaging evidence provides insights into how these networks contribute to language processing. Despite apparent similarities between mental operations involved in language comprehension (such as memory retrieval, prediction, and structure building) and those required by other cognitive domains, the MD network’s role in basic language comprehension appears more subtle than initially theorized^14–16^. By contrast, the MD network displays more robust engagement when language tasks introduce extra cognitive demands, such as retaining information in working memory or answering comprehension questions^17–19^. This can be understood as the language network implementing core computations for linguistic structure and meaning, while MD areas are recruited when task requirements demand increased cognitive control^15,20^. This dissociation is supported by evidence from multiple methodologies, including functional connectivity analyses^14,21^, studies of patients with focal brain lesions^22^, and investigations using naturalistic language stimuli^23,24^. These findings suggest that while the language network may implement certain computations on linguistic representations somewhat independently, it likely interacts with domain-general circuits in the MD network when tasks require additional cognitive control^20,25^.

While studies with adults have suggested relatively clear dissociation between language-specific and domain-general networks, documenting this dissociation in adults alone is insufficient for understanding the origin of uniquely human cognitive abilities. Several competing developmental trajectories could lead to the same adult organization: cortical regions might start as domain-general and gradually specialize through competitive interactions^4^, with language regions initially participating in broader cognitive functions like working memory before gaining linguistic selectivity. Alternatively, domain-general processes might bootstrap the development of language abilities^26–28^. A third pattern could be that there is an innate proto-organization of domain-specificity. Understanding how and when these networks dissociate during development is therefore critical for distinguishing between these possibilities and illuminating the origins of human cognitive architecture. Recent fMRI studies have demonstrated that this selective and specific language network is already established in children as young as 4 years of age, showing adult-like functional dissociation from domain-general cognitive processes^29,30^. However, due to the methodological constraints of fMRI, particularly the requirement for participants to remain still, the emergence and organization of these networks in even younger children remains largely unexplored. To the best of our knowledge, there is no existing exploration of the double dissociation of language and domain-general cortex in children younger than 4.

To address this gap, we employ functional near-infrared spectroscopy (fNIRS), which allows measurements while participants move relatively freely. fNIRS uses absorption of near-infrared light between emitters and detectors placed against the scalp to measure changes in oxygenated and deoxygenated hemoglobin concentrations that reflect neural activity^31^. This method has been successfully used to study early language processing in infants. Several studies have demonstrated that newborns already show specialized neural responses to speech in temporal and frontal regions^32–34^, while later work has suggested that language-specific responses continue to develop throughout infancy^35^. While these studies provide evidence for differential neural responses to speech versus reversed or degraded speech stimuli, they have not fully characterized the functional architecture of language processing in the developing brain. Specifically, showing greater activation to speech than non-speech does not establish whether these regions are specialized for language processing or serve more domain-general functions that happen to be engaged during speech processing. As Kanwisher^36^ emphasizes, in studies of functional specificity, demonstrating true specialization requires not only evidence of preferential responses to one category but should also test whether regions are exclusively engaged in processing that category. For example, the right temporoparietal junction (rTPJ) activates selectively when thinking about others’ thoughts, but not during closely related tasks like processing physical appearances or even other mental states such as hunger or pleasure. In additional to these conceptual challenges, mapping fNIRS measurement channels to specific cortical regions presents several methodological challenges, particularly in developmental populations. These challenges arise from anatomical variability in the relationship between external landmarks and underlying cortical structures, developmental variations in head size affecting measurement array coverage, and practical constraints in precise optode placement^37^. Moreover, even when measurement channels can be accurately mapped to specific anatomical regions, this anatomical localization does not guarantee functional correspondence. A given anatomical region often contains multiple functionally distinct sub-regions that vary in size and precise location across individuals^38,39^.

To overcome these methodological challenges, we employ a functional Channel of Interest (fCOI) approach for analyzing fNIRS data^37^. Unlike traditional approaches, which assume that measurement channels correspond to the same cortical regions across participants, the fCOI method identifies channels of interest by examining each individual’s functional activation patterns. This approach recognizes that the same channel position relative to scalp landmarks may measure different cortical regions across individuals due to variations in array placement, head size, and the relationship between surface landmarks and underlying cortical anatomy - a particularly important consideration in developmental research. The fCOI approach offers several key advantages for developmental studies^40^: it accounts for individual variability in brain anatomy and functional organization^41^, reduces noise from neighboring regions with different functional profiles, and provides a principled way to limit multiple comparisons while maintaining statistical power. This approach has been validated across multiple domains of cognitive development, including face processing and theory of mind^37,42,43^.

In the present study, we investigate the functional organization of language and domain-general networks in toddlers aged 2-4 years using fNIRS with the fCOI approach. Following Kanwisher’s^36^ framework for establishing functional specificity, our experimental design carefully contrasts responses across cognitive domains rather than simply identifying regions that respond preferentially to language. We focus on bilateral inferior frontal gyrus (IFG) regions given their well-documented heterogeneity in adult neuroimaging studies^13^. Specifically, prior research has established that Broca’s area within the left IFG contains two functionally distinct regions: a language-selective region responding specifically to linguistic input, and a domain-general region responding to diverse cognitive demands^13^. While anatomically adjacent, these regions belong to different networks - the left-lateralized frontotemporal language network and the bilateral frontoparietal multiple-demand (MD) network, respectively. Moreover, IFG’s anatomical characteristics^44^ (e.g. minimal hair interference, relatively thin skull in this region, proximity to cortical surface) maximize fNIRS signal quality. Therefore, IFG provides an optimal starting point for investigating whether specialized circuits emerge in core regions before extending to broader networks. Our experimental design includes a child-friendly language localizer adapted from Scott et al.^45^ that contrasts responses to auditorily presented engaging narrative clips versus acoustically degraded versions of the same stimuli. This localizer has been demonstrated to reliably identify language-selective regions across 45 languages from 12 language families, showing consistent activation patterns in the fronto-temporal language network regardless of the specific language being processed^46^. The passive listening nature of this task makes it particularly suitable for young children who may have difficulty with reading or cognitively demanding tasks with the confounding effects of task difficulty being controlled. The narrative stimuli can be easily customized with age-appropriate content to maintain children’s attention. For the cognitive control task, we used a spatial working memory task to engage the MD network for adults, as this paradigm has been consistently shown to be the most reliable and widely used task for localizing the MD network across numerous studies^10,22^. Using the same paradigm allows us to directly compare our fNIRS findings with the extensive fMRI literature. For children, we employed a Go/No-go task that probes inhibitory control - another key component of executive function that has been shown to engage the MD network^47^. While these tasks tap different aspects of executive function (working memory versus inhibitory control), both have been demonstrated to reliably activate the MD network and show similar patterns of increased activation under higher cognitive demands^9,10^. We implemented two conditions in the Go/No-go task: an easier condition with only "go" trials and a more demanding condition mixing "go" and "no-go" trials, paralleling the easy versus difficult manipulation in the adult working memory task. This modification was necessary as young children are unable to perform complex spatial working memory tasks^48^ but can engage with the more developmentally appropriate Go/No-go paradigm^49^ while still allowing us to probe domain-general cognitive control functions supported by the MD network. Through this approach, we aim to investigate whether these regions show distinct functional profiles for language processing versus domain-general cognitive demands in young children.

To validate our methodology, we first applied the individual fCOI approach to fNIRS data analysis in a sample of adults in experiment 1, ensuring that our source-detector array was capable of detecting activation from known functional regions in adult cortex. Then, in experiment 2, we examined whether the same language-preferring regions with functional profiles matching those established in the adult literature (i.e., selective responses to linguistic content and insensitivity to cognitive demands) could be observed in toddlers between 2-4 years of age using the same approach of identifying fCOIs in individual participants.

## Materials and Methods

### Experiment1

#### Participants

Twenty adults (between 18 and 26 years, 10 female) were recruited from the student population at Sichuan University. All participants had normal or corrected to normal vision, no history of neurological or psychiatric disorders, head trauma, seizures, or current use of psychoactive medications. None reported physical impairments that could affect task performance. All were right-handed and gave written informed consent. This study was approved by the Biomedical Ethics Review Committee of West China Hospital, Sichuan University (Protocol #2023-2376) and was conducted in accordance with the Declaration of Helsinki and CIOMS International Ethical Guidelines.

#### Procedure

Participants were seated in front of a 68 cm monitor at a distance of approximately 50 cm. Each participants completed (1) an auditory language localizer task^45^; (2) one spatial working memory task^6^.

In the language localizer task, participants listened to blocks of intact meaningful sentences and acoustically degraded speech following the procedure introduced in Scott et al.^45^. All the materials for this localizer were adopted to Mandarin from Gweon et al.^50^ and read by a native female speaker. We used female speakers because children tend to pay attention to female voices^51^. Thirty-two audio clips were created, consisting of 24 intact clips and 8 degraded clips (see the full list of stimuli in supplementary material). Each clip was 19-20 seconds in duration. Degraded speech clips were created from the intact versions using the procedure described below, resulting in muffled speech where the linguistic content was no longer intelligible. The degraded versions were created by first applying a low-pass filter (350 Hz pass-band) to the intact audio clips, then creating a noise track by randomizing the time-points of the original audio and multiplying it by the amplitude envelope of the intact clip. The noise track was low-pass filtered (8,000 Hz pass-band, 10,000 Hz stop frequency) to soften the highest frequencies and added to the filtered speech at a level that rendered it unintelligible. Each block lasted around 20 seconds. Each run included three blocks corresponding to three language conditions plus one block of degraded speech condition, with 10-second fixation blocks between blocks and additional fixation blocks at the beginning and end of the run. There are a total of eight runs. The task was originally designed to serve a dual purpose: localizing language processing and theory of mind networks. However, pilot testing revealed that children could not sustain this lengthy protocol, leading us to simplify the design to a single language condition for child participants in Experiment 2. For adult participants, we maintained all three language conditions but combined them into a single intact speech condition for analysis, contrasting it with the degraded speech condition. This resulted in an uneven distribution of stimuli between conditions (24 intact vs. 8 degraded clips). Participants were told that they would listen to some fun audio clips and some clips that were distorted in a way that makes it impossible to understand what the speaker is saying. They were instructed to listen attentively. Prior to the experiment, it was ensured that the volume level was sufficiently loud yet comfortable.

In the MD localizer task, participants performed a spatial working memory (WM) task involving a 3×4 grid. During each trial, squares appeared in different locations on the grid, either one at a time (easy condition) or two at a time (hard condition). Participants were required to remember these sequential locations. At the end of each trial, they were shown two sets of locations and had to select the set that matched the sequence they had just seen. They received feedback indicating whether their response was correct or incorrect^6^. Each trial followed a precise timing structure: beginning with a 500 ms fixation cross, followed by sequential grid presentations where each location was shown for 1000 ms. Participants then had up to 3750 ms to make their choice between the two location sets. After responding, they received feedback for 250 ms, followed by a fixation cross that filled the remaining time to ensure a consistent trial length (calculated as 4000 ms minus the response time minus 250 ms). Each run consisted of two blocks (one easy, one hard) with four trials per block, plus three 10-second fixation blocks interspersed at the beginning, middle, and end. Participants completed four runs in total. To ensure task comprehension, all participants completed a practice run before beginning the main experiment.

After each run the display paused until the participant initiated the next run by pressing a key on a keyboard located to his/her right. Participants were instructed to remain still and to focus on the screen throughout each run but were not asked to fixate on any point on the screen and could adjust their position between runs.

#### Data acquisition

fNIRS measurements were collected with a continuous wave system NirSmart (Danyang Huichuang Medical Equipment Co., Ltd., China) using wavelengths of 760 nm and 850 nm with a sampling rate of 11 Hz. 29 sources and 30 probes constituted 79 measurement channels of 3 cm (figure 1). The emitter and detector were placed according to the 10–20 system. The optodes were stabilized using a plastic holder and then affixed to participants’ heads over the frontal, temporal, occipital and parietal lobes in the left and right hemisphere using custom headgear. Prior to data collection, head circumference measurements were taken to ensure all participants fell within the suitable range for our standard cap size (54-58 cm). When wearing the optode cap, it was ensured that the Cz point of the electrode cap coincided with the Cz point of the scalp surface measurement. The spatial coordinates of sources, detectors, and anchor points (Nz, Cz, Al, Ar, lz, and other points from the international 10-20 system) were digitized using an electromagnetic 3D digitizer (Patriot, Polhemus, USA). These coordinates were then transformed into Montreal Neurological Institute (MNI) coordinates and mapped onto the MNI standard brain template using the spatial registration method in NirSpace (Danyang Huichuang Medical Equipment Co., Ltd, China).

**FIGURE 1:**
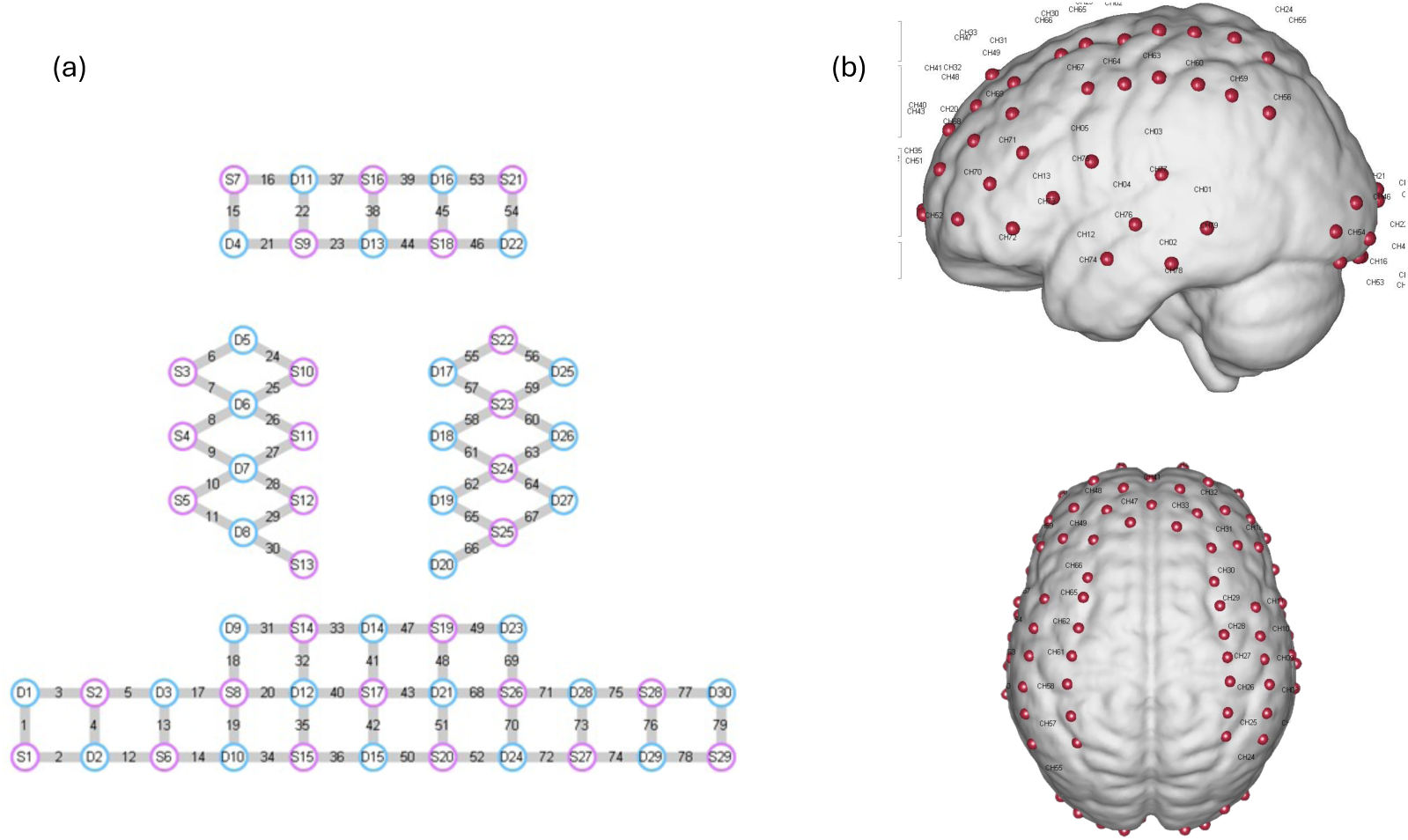
(a) Schematic representation of the fNIRS optode placement. Red circles indicate sources, blue circles indicate detectors, and yellow lines represent measurement channels. (b) Spatial registration result of the probe array. Red dots with number indicate channel positions on the MNI standard brain template.

#### Data preprocessing

Initial measurements of incident light reaching the detector for each channel were trimmed to remove excess time points from the beginning and end of data collection and were then preprocessed using the HomER NIRS processing package^52^. First, each run of the experiment was isolated from the continuous data and processed as a separate file to ensure that normalization of a given run was not skewed by more or less noisy portions of the data occurring at other times in the experiment^42^. Then, the negative intensity values resulting from noisy data were corrected using the hmrR_PreprocessIntensity_Negative function. Subsequently, channels were visually inspected and those lacking clear cardiac signal were manually excluded from individual participants’ data sets. Raw intensity measurements were then converted to optical density changes (ΔOD). Then a linear fit function was used to further remove the baseline drift. Next, a Temporal Derivative Distribution Repair (TDDR) was used to filter out motion artifacts, which can effectively remove baseline shift and spike artifacts without the need for any user-supplied parameters^53^ (Fishburn et al., 2019). The resulting time courses were then band-pass filtered with cut-off frequencies of 0.01–0.09 Hz to further remove slow drifts and high frequency and physiological noise^54^ (Pinti et al., 2019, butter filter 3 order for lowpass, 5 order for high pass). After that the ΔOD values were transformed into changes in oxyhemoglobin (HbO2) and deoxyhemoglobin (HHb) concentration according to the modified Beer-Lambert law (hmrOD2Conc, ppf = [1.0 1.0]). Then, the global components were removed using a principal component analysis spatial filter^55,56^(σ=46 deg; Zhang et al., 2016; Zhang et al., 2017), which is comparable to using short-source channels when the coverage of the measurement is much larger than the regions of interest^57^. And the concentration of each block is baseline corrected by 2s prior to the onset of stimuli.

#### Data analysis

We focused our analysis on changes in oxygenated hemoglobin concentration, as it has been demonstrated to be a more sensitive and reliable measure compared to deoxygenated or total hemoglobin concentration changes, which do not provide additional information beyond HbO2^58^. For each participant and channel, we computed the mean change in HbO2 concentration during each block, beginning 2 seconds post-stimulus onset to account for hemodynamic response lag. The analysis window extended to 25 seconds post-onset for the language localizer task and 38 seconds post-onset for the spatial working memory task, since different tasks’ blocks have different durations^58,59^. The differences in these responses across conditions were then analyzed using the fCOI approach.

First, we selected a set of channels ("search space")^37^ that plausibly covered regions of the scalp plausibly overlaying bilateral inferior frontal lobe across participants (see table 1 for probabilistic mapping of selected fNIRS channels to underlying cortical regions).

**TABLE 1.** Probabilistic mapping of fNIRS channels to underlying cortical regions showing Brodmann areas (based on MRIcro atlas) and corresponding MNI coordinates for each source-detector pair. Percentages indicate the probability of channel placement over the specified anatomical regions.

| Channel | MNI<br>coordinates | Anatomic label | Percentage | Search<br>Space |
| --- | --- | --- | --- | --- |
| **5** | 64.9643,<br>5.7288,<br>27.8566 | 4 - Primary Motor Cortex, 6 -<br>Pre-Motor and Supplementary<br>Motor Cortex, 43 - Subcentral<br>area, 44 - pars opercularis | 6.35%, 59.53%,<br>27.42%, 6.69% | RIFG |
| **13** | 61.2752,<br>22.4516,<br>12.4687 | 6 - Pre-Motor and Supplementary<br>Motor Cortex, 44 - pars<br>opercularis, 45 - pars triangularis,<br>48 - Retrosubicular area | 5.75%, 31.95%,<br>51.12%,<br>11.18% | RIFG |
| **14** | 57.049,<br>37.9202, -<br>2.4401 | 38 - Temporopolar area, 45 - pars<br>triangularis, 46 - Dorsolateral<br>prefrontal cortex, 47 - Inferior<br>prefrontal gyrus | 3.34%, 66.22%,<br>21.40%, 9.03% | RIFG |
| **17** | 53.2021,<br>33.3479,<br>29.3424 | 44 - pars opercularis, 45 - pars<br>triangularis, 46 - Dorsolateral<br>prefrontal cortex | 6.69%, 92.57%,<br>0.74% | RIFG |
| **19** | 49.1243,<br>47.253,<br>15.4105 | 45 - pars triangularis, 46 -<br>Dorsolateral prefrontal cortex | 48.70%,<br>51.30% | RIFG |
| **20** | 36.6861,<br>51.5968,<br>29.9235 | 9 - Dorsolateral prefrontal cortex,<br>45 - pars triangularis, 46 -<br>Dorsolateral prefrontal cortex | 4.17%, 3.70%,<br>92.13% | RIFG |
| **68** | -38.2423,<br>52.8662,<br>27.4047 | 45 - pars triangularis, 46 -<br>Dorsolateral prefrontal cortex | 12.45%,<br>87.55% | LIFG |
| **69** | -43.5347,<br>37.7257,<br>38.1227 | 9 - Dorsolateral prefrontal cortex,<br>44 - pars opercularis, 45 - pars<br>triangularis, 46 - Dorsolateral<br>prefrontal cortex | 25.45%, 4.55%,<br>39.09%,<br>30.91% | LIFG |
| **70** | -50.5051,<br>46.6936,<br>10.4438 | 45 - pars triangularis, 46 -<br>Dorsolateral prefrontal cortex | 53.45%,<br>46.55% | LIFG |
| **71** | -54.2983,<br>33.6329,<br>22.7694 | 45 - pars triangularis | 100% | LIFG |
| **72** | -56.3003,<br>37.7076, -<br>7.1488 | 38 - Temporopolar area, 45 - pars<br>triangularis, 46 - Dorsolateral<br>prefrontal cortex, 47 - Inferior<br>prefrontal gyrus | 8.14%, 48.06%,<br>19.77%,<br>24.03% | LIFG |
| **73** | -59.7137,<br>21.7772,<br>5.0031 | 6 - Pre-Motor and Supplementary<br>Motor Cortex, 38 - Temporopolar<br>area, 44 - pars opercularis, 45 -<br>pars triangularis, 48 -<br>Retrosubicular area | 4.03%, 15.44%,<br>14.09%,<br>36.24%,<br>30.20% | LIFG |
| 75 | -66.014, | 6 - Pre-Motor and Supplementary | 59.79%, | LIFG |
|  | 6.3358, | Motor Cortex, 43 - Subcentral | 31.12%, 4.90%, |  |
|  | 19.328 | area, 44 - pars opercularis, 48 - | 4.20% |  |
|  |  | Retrosubicular area |  |  |

The leave-one-run-out cross-validation procedure^42^ was implemented within each search space, which has demonstrated better sensitivity than other selection methods^40^. For each participant, one run was held out while the remaining runs were averaged. Within the predefined search space, we identified the eligible channel showing maximum contrast between intact speech and degraded speech for Language fCOI, and maximum contrast between hard and easy conditions for MD fCOI. This process was repeated iteratively for each run and participant. The identified channel was used as the fCOI definition to extract response values for each condition from each held-out run.

Our fCOI selection method does not guarantee the identification of reliably preferential channels in either search space. If none of the available channels recorded from cortex exhibited distinct responses between conditions, the selection process would just operate on random noise, resulting in no consistent response patterns in the independent data^37^. In other words, in cases where no robust preferential activation emerges during fCOI selection, any subsequent analyse may be driven by random variance rather than reflecting a true functional specialization, warranting cautious interpretation.

To compare the functional profile across the two networks, we also estimated the responses of the fCOIs to conditions that were not used to define them (i.e., hard and easy WM for the language fCOI, intact speech and degraded speech for the MD fCOI). For this analysis, we used all runs from the localizer task to define the fCOIs and all data from the other task to estimate their responses.

fCOIs were excluded from analyses if more than one-third of the channels in their search space were excluded^43^. Participants were excluded if they had fewer than 2 runs.

Our main analysis addressed three primary questions. First, regarding functional selectivity, we examined whether identified language fCOIs show selectivity to the properties of high-level language, specifically stronger activation in response to intact versus degraded speech^45^. Second, for functional specificity, we investigated whether language fCOIs exhibit specificity to language. A lack of difference in the language fCOI between the Easy and Hard conditions of the spatial working memory task, or a reverse pattern showing stronger responses to the Easy than the Hard condition, would suggest that the language network is not selectively engaged in this working memory task. Third, concerning domain-general processing, we needed to verify that the language fCOIs’ lack of response to cognitive load was not simply due to a failure of our tasks to engage domain-general function^60^. Therefore, we tested whether the MD fCOIs in our sample exhibited the expected selectivity and specificity for cognitive demands.

We employed a generalized linear mixed-effects model (GLMM) on individual trials to account for the unbalanced nature of the dataset (different numbers of runs across participants due to exclusion criteria) and individual differences in global signal strength. The analysis was implemented using MATLAB’s "fitglme" function with Maximum Penalized Likelihood (MPL) method. Our model structure was: HbO data ∼ condition + (1|subject) where condition represented the experimental conditions (intact vs degraded speech for language task; hard vs easy for cognitive/MD task). While inclusion of random slopes for condition by subject would be theoretically appropriate^61^, attempts to include these led to convergence failures. Separate models were run for two fCOI types in each search space (LIFG language, RIFG language, LIFG MD, RIFG MD) of two experiment contrasts (Intact > Degraded; Hard > Easy). To account for multiple comparisons, we applied Bonferroni correction separately for each task (language vs. cognitive control). In each task, the left and right IFG fCOIs formed a single family of two comparisons, yielding a family-wise threshold of α = 0.025 (0.05/2).

### Experiment 2

#### Participants

Thirty-five children between 2 and 4 years of age were recruited from preschools in Shanghai. Thirteen children were excluded because of failing to follow the experimental instructions. Twenty-two children (mean: 3 years 6 months; range: 2 years 5 months–4 years 11 months; 14 male, 8 female) were finally included in the study. Two children were aged 2 years 5 months to 2 years 11 months, seventeen were aged 3 years 0 months to 3 years 11 months, and three were aged 4 years 0 months to 4 years 11 months. According to parent reports, all participants had no developmental disorders, had normal or corrected-to-normal vision and hearing, and could understand basic instructions in Mandarin. Additionally, all children were full-term births (≥37 weeks gestation) with no significant medical complications at birth. Written informed consent was obtained from parents/legal guardians of all child participants prior to their inclusion in the study, in accordance with the requirements of the Biomedical Ethics Review Committee of West China Hospital, Sichuan University (Protocol #2023-2376). The study was conducted following the principles of the Declaration of Helsinki and CIOMS International Ethical Guidelines

#### Procedure

The procedure was similar to that of experiment 1 except that children under 3 years sat on their parents’ laps while viewing the display, and the experiment was discontinued, either during or between runs, if the participant became fussy, inattentive or if the parent indicated their wish to end the experiment. Each participants completed an auditory language localizer task and a Go/no-go task.

In the language localizer task, participants listened to blocks of intact meaningful sentences and acoustically degraded speech following the same procedure described in experiment 1. The materials were adapted from experiment 1 by selecting a subset of the adult stimuli based on a pre-experiment assessment of comprehension in a separate group of toddlers (n=3). While the adult version contained three language conditions per run, we selected only the material that showed the highest comprehension rates in our pre-experiment (accurate responses to simple comprehension questions. E.g. "Was the girl helping her father?" or "Was the rabbit in the garden?"). The selected narratives were shortened from the original 20-second duration to 18-second segments to accommodate shorter attention spans, while maintaining the core narrative structure. Each block lasted 18 seconds, with 6-second fixation blocks between blocks and additional fixation blocks at the beginning and end of the run. Each run included one language condition block plus one degraded speech condition block. There are four total runs. As in experiment 1, participants were told that they would listen to some fun audio clips and some clips that were distorted in a way that makes it impossible to understand what the speaker is saying. They were instructed to listen attentively. Prior to the experiment, it was ensured that the volume level was sufficiently loud yet comfortable.

In the MD localizer task, participants completed a Go/no-go task. A training session before the experiment was conducted to ensure task comprehension. The training included practice trials for both "go" (simple response) and "go/no-go" (response inhibition) conditions with performance feedback, continuing until participants demonstrated reliable response patterns. The formal experiment consisted of four runs, each containing alternating "go" and "go/no-go" blocks separated by 10-second fixation periods. Each block began with a 3-second instruction screen followed by 12 trials lasting 12 seconds total. In "go" blocks, participants pressed a key with their right index finger in response to both cat and dog images. In "go/no-go" blocks, participants pressed the key for chicken images (go) but withheld responses for duck images (no-go). A study personnel monitored participant performance during scanning.

#### Data acquisition

fNIRS measurements were collected with a continuous wave system NirSmart (Danyang Huichuang Medical Equipment Co., Ltd., China) using wavelengths of 730nm, 808 nm and 850 nm with a sampling rate of 11 Hz. 26 sources and 27 probes constituted 81 measurement channels of 3 cm (see figure 2). The optodes were stabilized using a plastic holder and then affixed to participants’ heads over the frontal, temporal, occipital and parietal lobes in the left and right hemisphere using custom headgear. When wearing the optode cap, it was ensured that the Cz point of the electrode cap coincided with the Cz point of the scalp surface measurement.

**FIGURE 2.**
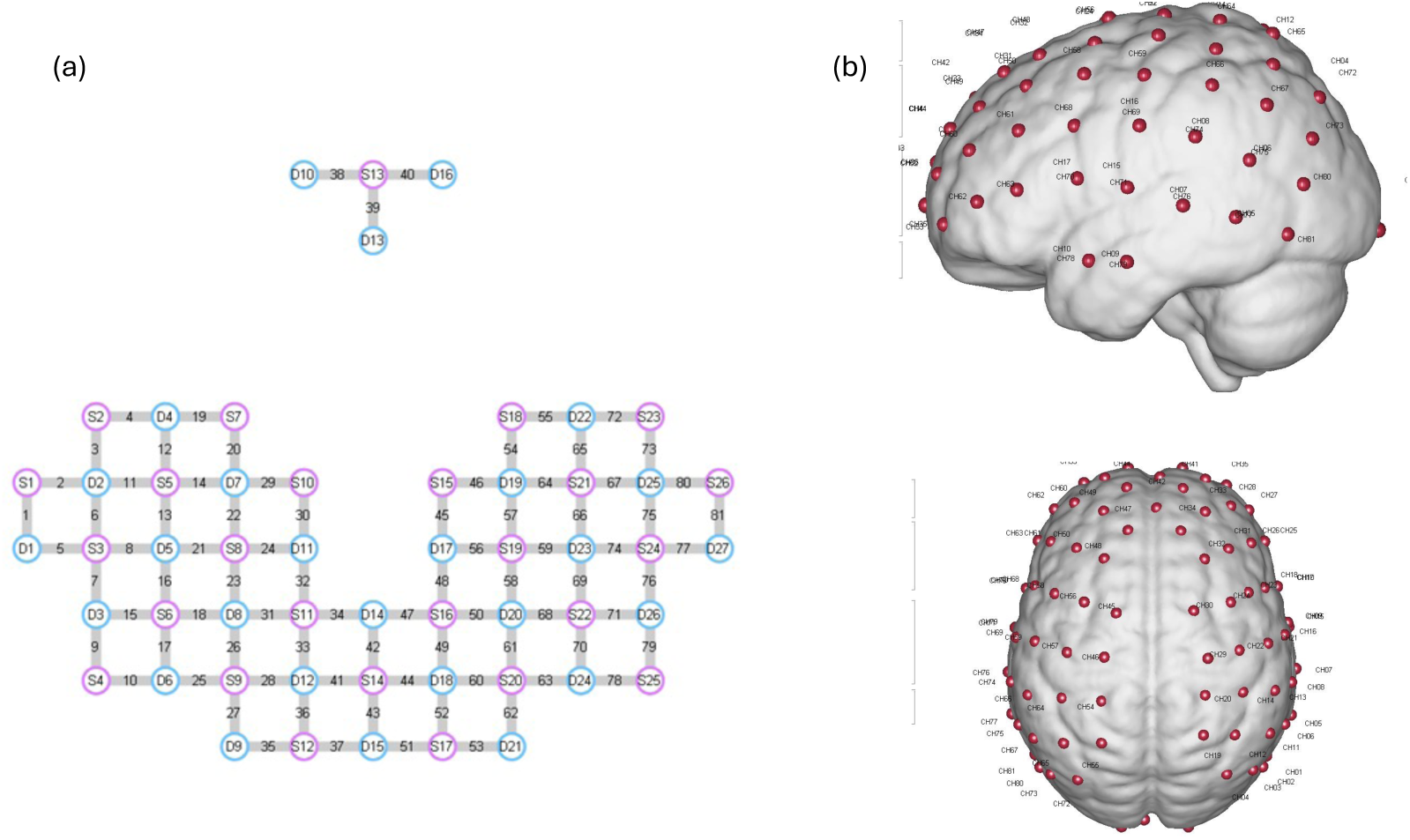
: (a) Schematic representation of the fNIRS optode placement. Red circles indicate sources, blue circles indicate detectors, and yellow lines represent measurement channels. (b) Spatial registration result of the probe array. Red dots with number indicate channel positions on the MNI standard brain template.

#### Data preprocessing

Data were preprocessed in the same manner as Experiment 1, with the exception of the partial pathlength factor (ppf) in the modified Beer-Lambert law calculation. Given the different optical properties of toddler brain tissue compared to adults, we used a ppf value of [2.0 2.0 2.0], which better accounts for the light scattering and absorption characteristics in infant participants^62^.

#### Data analysis

For each participant and channel, mean HbO2 concentration changes were calculated per trial. The analysis window started 4 seconds after stimulus onset and lasted until 17 seconds post-onset for the language localizer task or 12 seconds post-onset for the Go/no-go task. This timing accounts for the HRF lag time commonly assumed in children’s NIRS research^63,64^. Language and MD selective regions were identified and mean HbO responses were extracted using individual fCOI approaches within the same search space (see table 2 for probabilistic mapping of selected fNIRS channels to underlying cortical regions), following Experiment 1’s methodology. Inclusion criteria required children to complete at least two runs per experiment, ensuring sufficient trials for the leave-one-run-out procedure. Additional exploratory analyses incorporating age as a fixed effect in the generalized linear mixed models are reported in the supplementary material.

**TABLE 2.**
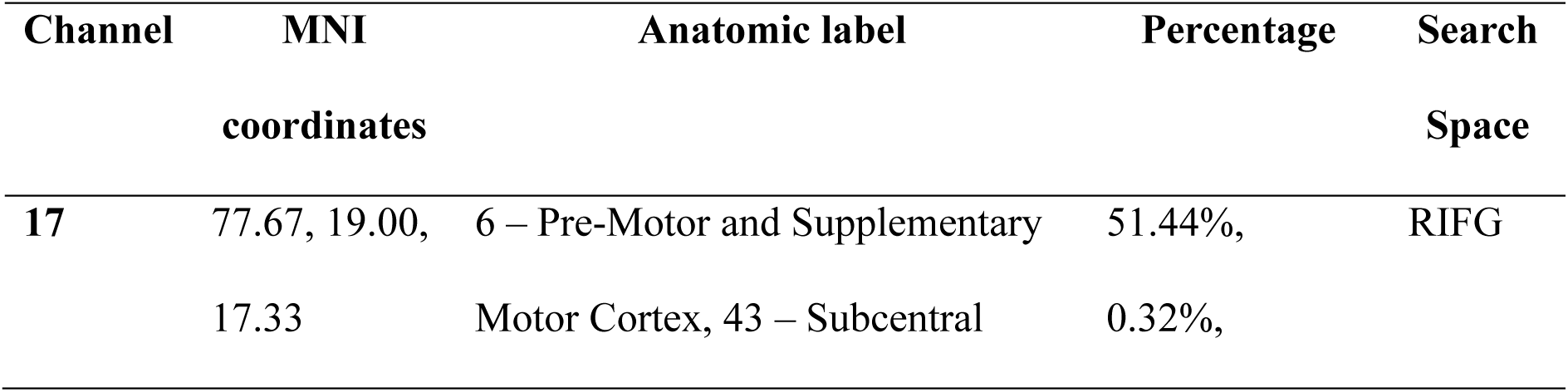
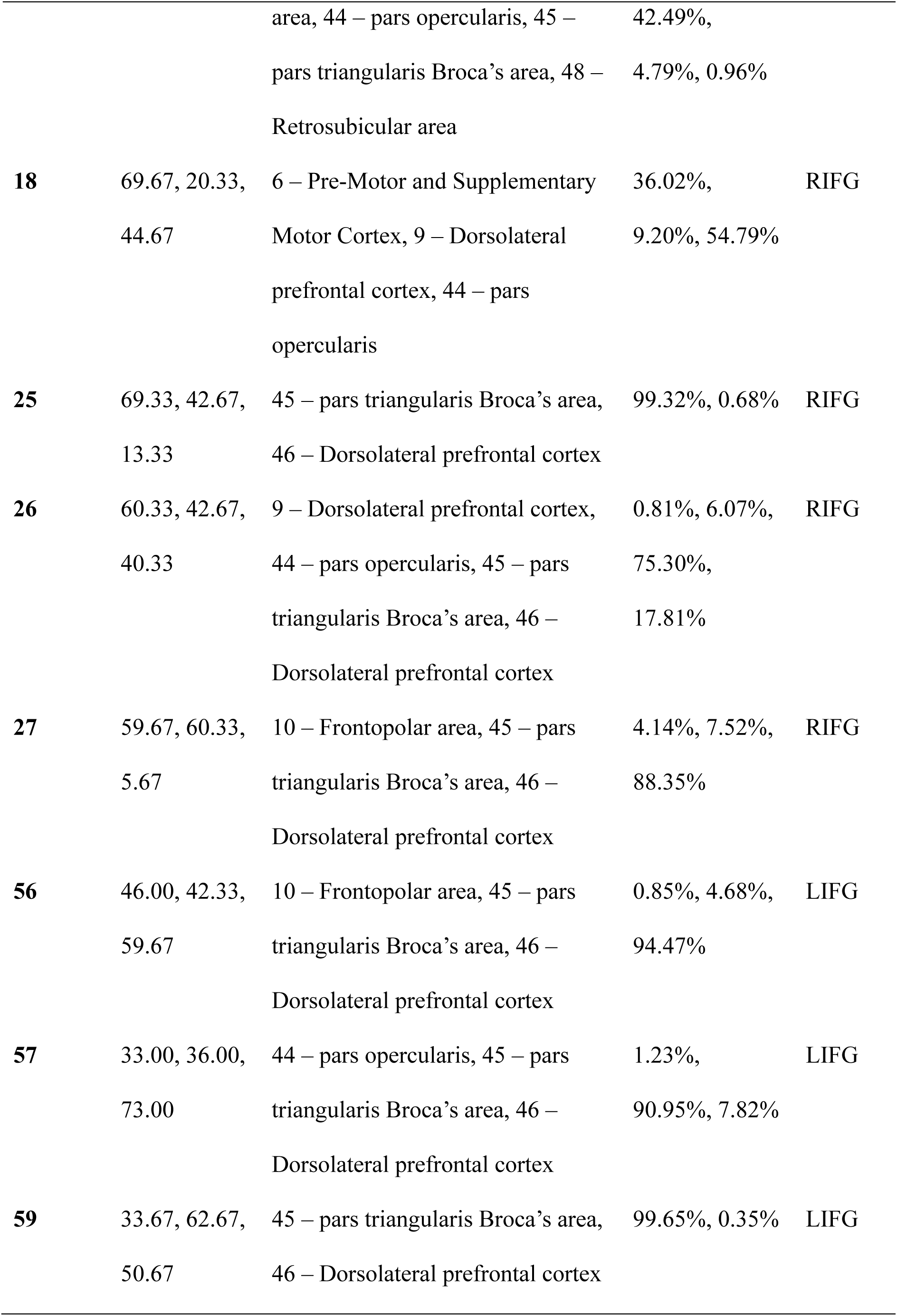
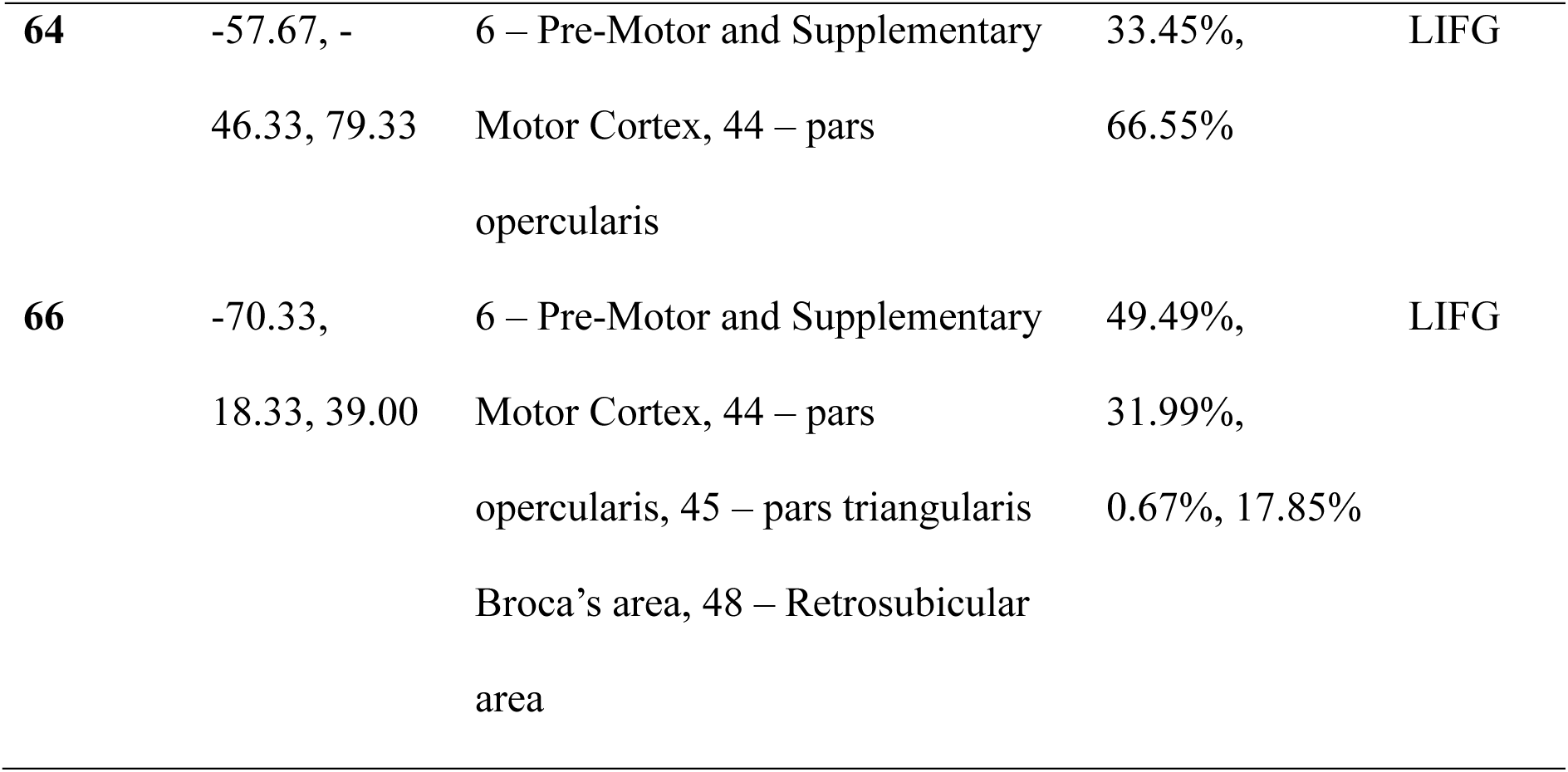
Probabilistic mapping of fNIRS channels to underlying cortical regions showing Brodmann areas (based on MRIcro atlas) and corresponding MNI coordinates for each source-detector pair. Percentages indicate the probability of channel placement over the specified anatomical regions.

| Channel | MNI<br>coordinates | Anatomic label | Percentage | Search<br>Space |
| --- | --- | --- | --- | --- |
| 17 | 77.67, 19.00,<br>17.33 | 6 – Pre-Motor and Supplementary<br>Motor Cortex, 43 – Subcentral | 51.44%,<br>0.32%, | RIFG |
|  |  | area, 44 – pars opercularis, 45 – | 42.49%, |  |
|  |  | pars triangularis Broca's area, 48 – | 4.79%, 0.96% |  |
|  |  | Retrosubicular area |  |  |
| **18** | 69.67, 20.33, | 6 – Pre-Motor and Supplementary | 36.02%, | RIFG |
|  | 44.67 | Motor Cortex, 9 – Dorsolateral | 9.20%, 54.79% |  |
|  |  | prefrontal cortex, 44 – pars |  |  |
|  |  | opercularis |  |  |
| **25** | 69.33, 42.67, | 45 – pars triangularis Broca's area, | 99.32%, 0.68% | RIFG |
|  | 13.33 | 46 – Dorsolateral prefrontal cortex |  |  |
| **26** | 60.33, 42.67, | 9 – Dorsolateral prefrontal cortex, | 0.81%, 6.07%, | RIFG |
|  | 40.33 | 44 – pars opercularis, 45 – pars | 75.30%, |  |
|  |  | triangularis Broca's area, 46 – | 17.81% |  |
|  |  | Dorsolateral prefrontal cortex |  |  |
| **27** | 59.67, 60.33, | 10 – Frontopolar area, 45 – pars | 4.14%, 7.52%, | RIFG |
|  | 5.67 | triangularis Broca's area, 46 – | 88.35% |  |
|  |  | Dorsolateral prefrontal cortex |  |  |
| **56** | 46.00, 42.33, | 10 – Frontopolar area, 45 – pars | 0.85%, 4.68%, | LIFG |
|  | 59.67 | triangularis Broca's area, 46 – | 94.47% |  |
|  |  | Dorsolateral prefrontal cortex |  |  |
| **57** | 33.00, 36.00, | 44 – pars opercularis, 45 – pars | 1.23%, | LIFG |
|  | 73.00 | triangularis Broca's area, 46 – | 90.95%, 7.82% |  |
|  |  | Dorsolateral prefrontal cortex |  |  |
| **59** | 33.67, 62.67, | 45 – pars triangularis Broca's area, | 99.65%, 0.35% | LIFG |
|  | 50.67 | 46 – Dorsolateral prefrontal cortex |  |  |
| **64** | -57.67, - | 6 – Pre-Motor and Supplementary | 33.45%, | LIFG |
|  | 46.33, 79.33 | Motor Cortex, 44 – pars opercularis | 66.55% |  |
| **66** | -70.33, | 6 – Pre-Motor and Supplementary | 49.49%, | LIFG |
|  | 18.33, 39.00 | Motor Cortex, 44 – pars opercularis, 45 – pars triangularis | 31.99%, |  |
|  |  | Broca’s area, 48 – Retrosubicular area | 0.67%, 17.85% |  |

## Results

### Experiment 1

No participants were excluded based on our criteria for channel rejection and fCOI rejection. For the subsequent analyses, we included all data after removing the excluded noisy channel and any runs where noisy channels comprised more than one-third of the search space. Table 3 provides a summary of the available runs across participants for different fCOI types (See the distribution of identified channel for each type of fCOI in supplementary material in figure S4-S8).

**TABLE 3.** Summary of the available runs across participants for different fCOI types.

| **fCOI** | **Total LAN Subjects** | **Mean LAN Runs** | **Total MD** | **Mean MD** |
| --- | --- | --- | --- | --- |
| **Type** | **Included** | **(SDs)** | **Subjects** | **Runs (SDs)** |
|  |  |  | **Included** |  |
| **LAN** | 20 | 6.95(0.683) | 20 | 3.85(0.489) |
| **LIFG** |  |  |  |  |
| **LAN** | 20 | 7.15(0.3663) | 20 | 3.85(0.489) |
| **RIFG** |  |  |  |  |
| **MD** | 20 | 7.10(0.4472) | 20 | 3.90(0.4472) |
| **LIFG** |  |  |  |  |
| **MD** | 20 | 7.15(0.3663) | 20 | 3.90(0.4472) |
| **RIFG** |  |  |  |  |

The language fCOIs showed clear selectivity for language processing. In both hemispheres, these channels demonstrated significantly stronger responses to intact versus degraded speech. The LIFG language fCOI showed a significant effect of condition (β = −8.92e-06, SE = 3.04e-06, t = −2.9365, p = 0.0036), with higher activation for intact speech. Similarly, the right inferior frontal gyrus (RIFG) language fCOI exhibited a significant preference for intact speech (β = −1.18e-05, SE = 3.36e-06, t = −3.5248, p = 0.00049). Importantly, when tested on the spatial working memory task, neither the left nor right language fCOIs showed significant modulation by cognitive demand (LIFG: β = −4.66e-06, SE = 5.31e-06, t = −0.87838, p = 0.381; RIFG: β = −2.1894e-06, SE = 4.9866e-06, t = −0.43906, p = 0.661). This pattern suggests that the language fCOIs are specifically selective for language processing and do not respond systematically to increased cognitive load in non-linguistic tasks (figure 3a & 3b).

**FIGURE 3.**
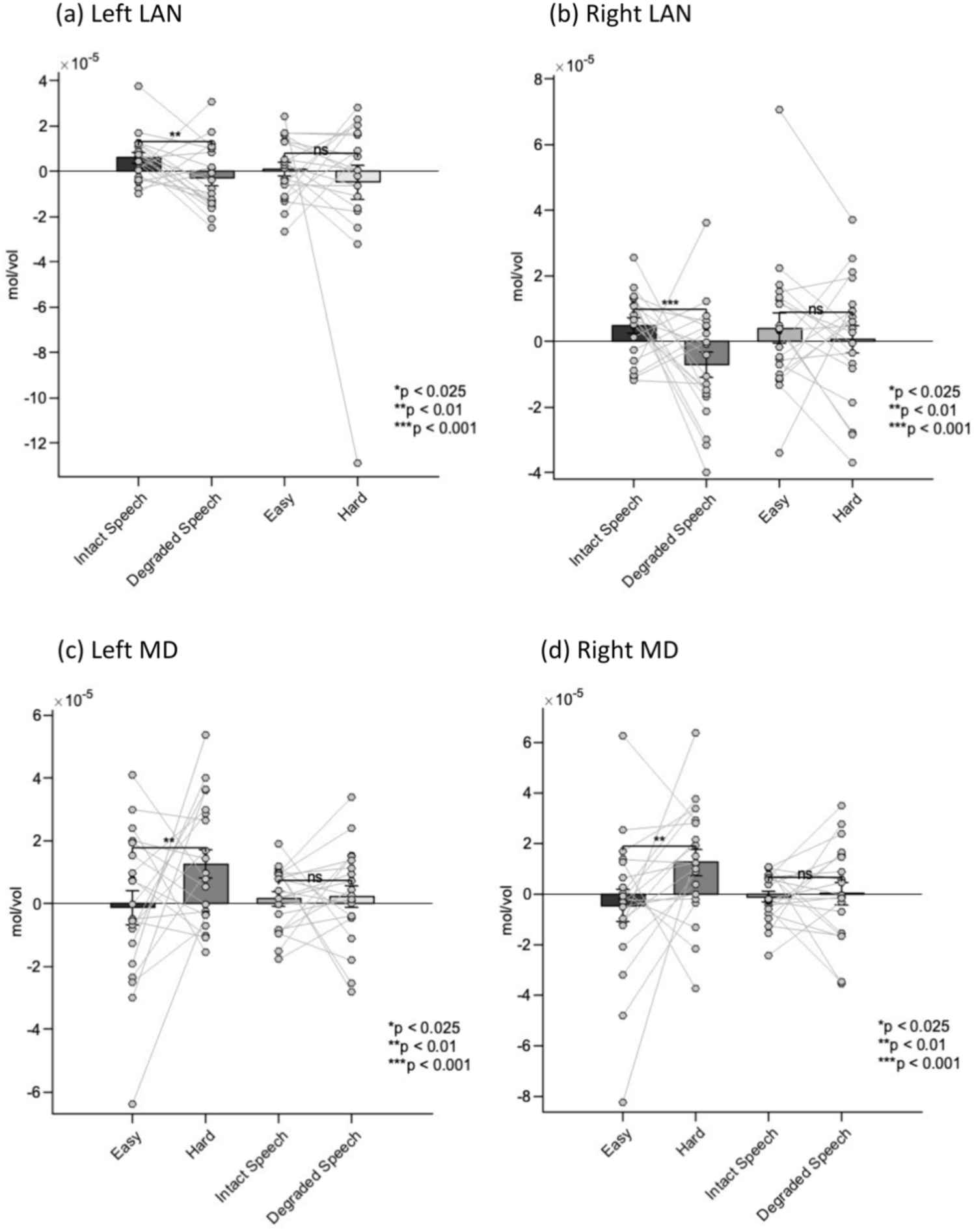
Results from Experiment 1: Mean HbO2 responses in the selected fCOIs (a) Left hemisphere language fCOI responses showing significant selectivity for intact versus degraded speech (p = 0.0036) and no significant modulation by cognitive demand (p = 0.381). (b) Right hemisphere language fCOI responses showing significant selectivity for intact versus degraded speech (p = 0.00049) and no significant modulation by cognitive demand (p = 0.661). (c) Left hemisphere MD fCOI responses showing significant sensitivity to cognitive demand (p = 0.0095) and no significant difference between speech conditions (p = 0.988). (d) Right hemisphere MD fCOI responses showing significant sensitivity to cognitive demand (p = 0.0011) and no significant difference between speech conditions (p = 0.756). Error bars represent standard error of the mean.

The MD fCOIs demonstrated a complementary pattern of activation. Both left and right MD fCOIs showed significant sensitivity to cognitive demand in the spatial working memory task. The LIFG MD fCOI exhibited stronger activation for harder versus easier conditions (β = 1.34e-05, SE = 5.09e-06, t = 2.6279, p = 0.0095), as did the RIFG MD fCOI (β = 1.75e-05, SE = 5.26e-06, t = 3.3278, p = 0.0011). In contrast, when tested on the language task, neither MD fCOI showed significant differentiation between intact and degraded speech (LIFG: β = 5.04e-08, SE = 3.25e-06, t = 0.015523, p = 0.988; RIFG: β = 1.01e-06, SE = 3.25e-06, t = 0.31183, p = 0.756). This finding indicates that the MD network responds selectively to cognitive demand but not to linguistic properties per se (figure 3c & 3d).

These results demonstrate a double dissociation between the language and MD networks, with language fCOIs showing selectivity for linguistic processing but not cognitive demand, and MD fCOIs showing the opposite pattern.

### Experiment 2

Table 4 provides a summary of the included participants and available runs across participants for different fCOI types based on the criteria for fCOIs rejection we described in experiment 1 (See the distribution of identified channel for each type of fCOI in supplementary material, Figure S1-S4).

**TABLE 4.** Summary of the available runs across participants for different fCOI types.

| **fCOI Type** | **Total LAN Subjects Included** | **Mean LAN Runs (SDs)** | **Total MD Subjects Included** | **Mean MD Runs (SDs)** |
| --- | --- | --- | --- | --- |
| **LAN** | 22 | 3.54(0.510) | 21 | 2.24(0.436) |
| **LIFG** |  |  |  |  |
| **LAN** | 22 | 3.50(0.597) | 17 | 2.23(0.437) |
| **RIFG** |  |  |  |  |
| **MD** | 22 | 3.50(0.597) | 22 | 2.27(0.457) |
| **RIFG** |  |  |  |  |
| **MD** | 21 | 3.38(0.74) | 21 | 2.24(0.539) |
| **LIFG** |  |  |  |  |

Analysis of the language fCOIs in children showed a partially overlapping pattern with adults. The LIFG language fCOI maintained significant selectivity for language processing, showing stronger responses to intact versus degraded speech (β = −5.21e-07, SE = 1.69e-07, t = −3.0781, p = 0.0025). However, unlike adults, the RIFG language fCOI did not show significant differentiation between intact and degraded speech (β = −7.49e-08, SE = 1.90e-07, t = 0.1211, p = 0.693). Consistent with the adult findings, neither language fCOI showed significant modulation by cognitive demand in the Go/no-go task (LIFG: β = −3.22e-08, SE = 1.46e-07, t = −0.21987, p = 0.826; RIFG: β = −8.69e-08, SE = 2.01e-07, t = −0.72473, p = 0.667). This suggests that even in children, the language fCOI we identified maintains specificity for language processing over general cognitive demands (figure 4a & 4b).

**FIGURE 4.**
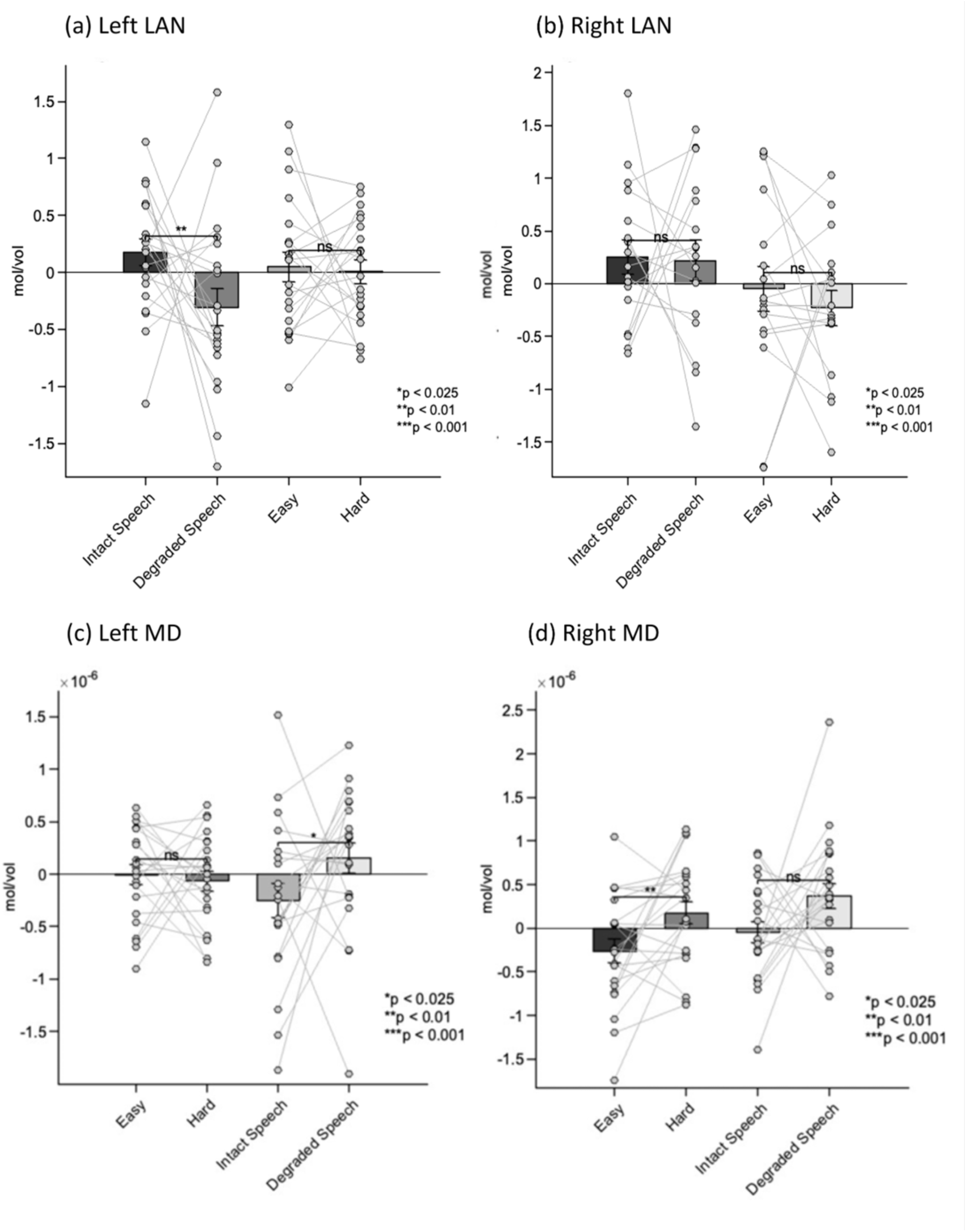
Results from Experiment 2: Mean HbO2 responses in the selected fCOIs (a) Left hemisphere language fCOI responses showing significant selectivity for intact versus degraded speech (p = 0.0025) and no significant modulation by cognitive demand (p = 0.826). (b) Right hemisphere language fCOI responses showing no significant selectivity for speech conditions (p = 0.693) and no significant modulation by cognitive demand (p = 0.667). (c) Left hemisphere MD fCOI responses showing no significant sensitivity to cognitive demand (p = 0.4917) but significant preference for degraded speech (p = 0.015). (d) Right hemisphere MD fCOI responses showing significant sensitivity to cognitive demand (p = 0.00323) and marginal preference for degraded speech (p = 0.046). Error bars represent standard error of the mean.

The MD fCOI we identified in children showed a more complex pattern than expected. Behavioral performance confirmed the task difficulty manipulation, with participants showing higher accuracy on go trials (M = 81%, SD = 12%) compared to no-go trials (M = 73%, SD = 15%; t(21) = 2.25, p = 0.035). Only the right MD fCOI displayed higher sensitivity to cognitive demand in the Go/no-go task (significantly stronger responses to the harder versus easier condition; RIFG: β = 4.49e-07, SE = 1.48e-07, t = 3.0185, p = 0.00323). By contrast, the left MD fCOI did not show a significant difference between these conditions (LIFG: β = −8.63e-08, SE = 1.25e-07, t = −0.69016, p = 0.4917). When we tested these fCOI in the language task, an unexpected pattern emerged: although the left MD fCOI was not strongly modulated by cognitive control demands, it nonetheless responded more robustly to degraded than to intact speech (β = 4.2037e-07, SE = 1.7142e-07, t = 2.4523, p = 0.015313). The right MD fCOI showed a similar trend though not reaching statistical significance after correction for multiple comparisons (β = 3.6637e-07, SE = 1.8245e-07, t = 2.0081, p = 0.046). These findings suggest that only the right MD fCOI displays the characteristic MD network profile of responding to cognitive control demands but not to linguistic properties. The left MD fCOI’s lack of modulation by task difficulty suggests that an adult-like MD network component may not yet be established in the left IFG in our sample, warranting caution in interpreting its preferential response to degraded speech as we have discussed in the method part (figure 4c & 4d).

## Discussion

Our study investigates how the brain supports language processing in early childhood, focusing on whether neural circuits are specialized for language or share with other cognitive domains. By employing fNIRS with an individual functional channel of interest approach, we assessed both adults and toddlers on language and cognitive control tasks to seek evidence for distinct functional profiles in the left and right inferior frontal regions. Our adult findings validated the fCOI method by replicating well-established dissociations between language-specific and domain-general processor, providing a foundation for investigating the developmental period. Our toddler results suggest that left-hemisphere language selectivity emerges early and remains robust, while right-hemisphere organization shows signs of ongoing maturation.

In line with adult findings and recent work in older children^29,65^, toddlers demonstrated robust language selectivity in the left IFG. The language fCOI within this area showed specific responses to intact versus degraded speech, mirroring the adult pattern, while remaining insensitive to the manipulation of cognitive demands in the Go/No-Go task. This supports the hypothesis of early domain specificity in language processing, as the language selective region within left IFG region exhibited both selectivity and specificity for language^36,60^. Our toddler data align with evidence that even very young children recruit the left frontal cortex for high-level linguistic processing^33,35,66^. Moreover, our findings revealed a developmental asymmetry in language processing between hemispheres. While our adult participants showed bilateral IFG fCOIs with selectivity for meaningful speech, toddlers did not exhibit statistically reliable language responses in the RIFG language fCOI. This asymmetry aligns with previous developmental research and recent findings in older children. Hiersche et al.^29^, using a similar speech localizer task comparing sentences to acoustically degraded speech, also found no significant language selectivity in the right IFG. Early in development, infant studies have documented limited right-hemisphere sensitivity to language in speech versus control conditions^33,66^. A developmental shift occurs later: children ages 5-11 show right hemisphere engagement, particularly during complex linguistic tasks requiring greater syntactic processing or elevated linguistic demands^67–69^. Our observation of strong LIFG but nonsignificant RIFG selectivity in toddlers (ages 2-4) suggests that RIFG language specialization follows a prolonged developmental trajectory. While LIFG supports core linguistic computations from early development, RIFG may serve complementary functions that become increasingly important as language abilities mature. Studies suggest RIFG involvement in prosodic processing, pragmatic aspects of language, and discourse-level comprehension^70,71^. The absence of RIFG language selectivity in toddlers suggest that these higher-level linguistic functions develop later, potentially aligning with children’s growing ability to process complex narrative^72^ and social aspects of language^73^. These findings support a developmental model where left-hemisphere language selectivity emerges early and remains stable, while right-hemisphere language involvement develops gradually throughout childhood^67,74^.

With regard to domain-general processing, our findings revealed a distinct pattern in toddlers from adults: only the right MD fCOI showed selective sensitivity to increased cognitive demands during the Go/No-Go task. These MD fCOIs demonstrated functional profiles that contrasted with language fCOIs - while language regions responded selectively to intact versus degraded speech but showed no sensitivity to cognitive demands, MD fCOIs exhibited the opposite pattern. Importantly, only the right MD fCOI displayed the characteristic MD network profile of responding to cognitive control demands but not to linguistic properties. This toddler-specific pattern diverges from established findings in both adults and older children (approximately 7-12 years). Adults show bilateral MD network responses to escalating task difficulty, and older children demonstrate bilateral prefrontal recruitment during inhibitory control tasks^75–78^. The developmental differences likely reflect the ongoing structural and functional specialization of the MD network in early childhood. Our findings of right MD responsiveness in toddlers, coupled with previous fNIRS and fMRI studies, suggest that very young children’s prefrontal cortex involvement in executive tasks may initially manifests in right hemisphere circuits compared to older groups^79,80^. This aligns with evidence of protracted maturation for left frontal functions^75,77,81^. The left MD fCOI’s lack of modulation by task difficulty suggests that an adult-like MD network component may not yet be established in the left IFG in our sample. This developmental pattern is further corroborated by recent work from Schettini et al.^30^ who similarly found that the left IFG in children did not show significant selectivity for cognitive demand. While the left MD fCOI exhibited stronger activation to degraded compared to intact speech, this pattern should be interpreted cautiously, as it lacks the hallmark feature of the MD network—sensitivity to cognitive demand—and may reflect immature functional engagement in this region. An alternative possibility is that degraded speech, which is more challenging to understand than intact speech, elicits more effortful processing in children, thereby driving up activity in what would eventually become a domain-general region. In other words, although the left MD fCOI does not show the classic adult-like sensitivity to cognitive demands, it may nonetheless be recruited when children strain to extract meaning from ambiguous input. Together, these findings support previous research indicating ongoing development of prefrontal involvement in domain-general control^30,74,76^, with bilateral integration emerging later in development as children increasingly recruit both hemispheres for executive tasks.

Our use of the fCOI approach addressed several methodological challenges in fNIRS studies. First, it circumvents the limitations of the ‘reverse inference’ problem which plagues many fNIRS studies that rely on inferring cognitive functions solely from anatomical correspondence with prior findings. This practice is problematic as the same anatomical location may support different functions across individuals^82^, particularly in developmental research where variations in head size, brain organization, and probe placement complicate anatomical localization^41^. Our fCOI approach addresses these limitations by identifying regions based on functional properties rather than assumed anatomical locations, effectively accommodating individual variability in neural functional anatomy across development. Second, it provided enhanced statistical power while controlling for multiple comparisons, which is crucial when working with limited developmental data^40^. Third, the implementation of the same localizer task used by fMRI studies^7,29,65^ allowed for direct comparison of properties of specific functional regions over developmental stages and imaging modalities. Our successful identification of functionally distinct networks in both adults and toddlers demonstrates the robustness of this approach across developmental stages.

Our experimental design also addressed several methodological challenges. The implementation of a leave-one-run-out cross-validation procedure for fCOI selection avoided the "double dipping" issue in neuroimaging studies^83^, enhancing the reliability of our findings. By employing passive language tasks contrasting meaningful versus degraded speech, we controlled for confounding effects of task difficulty while maintaining prosodic and rhythmic features^45^. This paradigm, extensively validated in adult studies across different languages, allowed us to specifically probe the processing of high-level linguistic features such as semantics and syntax^46^. The careful selection of age-appropriate tasks for probing the MD network—spatial working memory for adults and Go/No-go for children—enabled us to examine domain-general processing while accommodating developmental constraints. By comparing responses to both linguistic and non-linguistic tasks within the same participants, we directly assessed the specificity of these networks to language processing.

Despite these methodological strengths, several limitations warrant discussion. First, like most fNIRS experiments, our study faced challenges of limited and non-continuous cortical coverage. While our probe arrangement provided good coverage of bilateral inferior frontal regions, some cortical areas may have fallen into measurement "blind spots"^84^. Second, we did not have access to short-separation channels in our study. Short-separation channels are efficient in accounting for and removing superficial hemodynamic signals originating from the scalp and skull, thereby isolating the cortical activity of interest^85,86^. The absence of short-separation channels may have led to contamination of our signals by systemic physiological artifacts. Although we have applied global regression method to filter out systemic physiological artifacts^55^, systemic physiological artifacts may still potentially affect the specificity and accuracy of our findings. Third, different cognitive tasks were used for adults and children. While both tasks engage the MD network, the cognitive demands they place on participants differ. However, this adaptation was necessary given the age-appropriate task requirements, and both tasks share the feature of manipulating cognitive control demands. Fourth, while our peak-channel selection approach within predefined anatomical regions of interest was optimized for our specific hypotheses, alternative approaches might be valuable for different research questions. Additionally, we did not examine behavioral metrics of language development, which limits our ability to differentiate between changes in language selectivity due to maturation versus language skill development^87^. Finally, our focus was primarily on the inferior frontal cortex, which constrained our ability to fully characterize both the MD network and the language-selective network. While the IFG is a critical hub for domain-general cognitive control^9^ and language processing^10^, the broader language network encompasses distributed temporal, parietal, and superior frontal regions. By restricting our measurements to the IFG, we may have overlooked crucial activity in other regions of the language network (e.g., the posterior temporal lobe) that could further elucidate the transition from domain-general to domain-specific processes in language development. Consequently, caution is warranted when generalizing our findings to the broader MD and language networks.

Future research should extend these findings in two main ways. First, longitudinal studies beginning at earlier ages^5,66^ are needed to clarify whether language networks transition from domain-general to domain-specific functioning over time. Second, studies should examine neural specialization at key language acquisition milestones by collecting brain measures when children reach specific linguistic achievements (e.g., first words, word combinations, complex sentences). This milestone-based approach^7,87^ would illuminate how maturing brain circuits support children’s emerging linguistic abilities.

To conclude, using the methodologically improved fCOI approach with fNIRS, we replicated well-established patterns of bilateral selectivity and specificity in adult participants during linguistic and cognitive control tasks. In toddlers, we found early specialization of the language-selective component within LIFG, while the homologous right hemisphere region had not yet developed language selectivity. On cognitive control tasks, toddlers showed adult-like patterns in the within RIFG only. Notably, the selected MD fCOI in toddler’s LIFG only showed engagement with degraded speech rather than the broader domain-general profile seen in adults, suggesting ongoing maturation of domain-general circuits. These findings advance our understanding of how specialized brain circuits for language emerge during early childhood and establish the utility of the fCOI approach for studying functional organization in the developing brain.

## Code and Data Availability

The data that support the findings of this study are available from the corresponding author upon reasonable request. The data are not publicly available due to privacy or ethical restrictions involving data from minor participants.

## Acknowledgments

We express our sincere gratitude to all participating families and children for their invaluable contributions to this study. We thank WEN Jian for her dedicated efforts in data collection. We are deeply grateful to Evelina Fedorenko, Arturo E. Hernandez, CHEN Xiaocong, and QI Zhenghan for their insightful discussions and guidance. Our thanks extend to CHEN Jiahan, YANG Jingdan, and GENG Luyuan for their assistance in adapting the materials into Chinese. We also acknowledge the crucial support of the preschools affiliated with the Shanghai Maternal and Child Health Center in participant recruitment.

This research was supported by The Hong Kong Polytechnic University Presidential PhD Fellowship Scheme (PPPFS) and Ordos Clinical Medicine Translational Innovation Center Program Science and Technology Cooperation Program of Shanghai Jiaotong University in Inner Mongolia Autonomous Region—Action Plan of Shanghai Jiaotong University for "Revitalizing Inner Mongolia through Science and Technology" (KJXM2024-11-01).

## Biography

Haolun Luo is a PhD student at the Hong Kong Polytechnic University. He received his BS degree in Hearing and Speech Rehabilitation from Sichuan University in 2022. His research interests focus on using fNIRS to assess children’s cognitive and language abilities, with the goal of developing objective biomarkers for child development.

Tao Yu is an Assistant Professor at the Bio-X Institutes, Shanghai Jiao Tong University, and a PI at Shanghai Mental Health Center. He received his PhD from Sichuan University and completed postdoctoral research at Shanghai Jiao Tong University and King’s College London. His research includes work on precision medicine, chronic non-communicable disease prevention, and cognitive neuroscience.

Qun Li received her PhD from The Chinese University of Hong Kong. She is recognized as a high-level overseas talent in Sichuan Province. Her research focuses on language assessment, child language acquisition, language and cognition in children with language disorders, and speech rehabilitation for hearing-impaired children. She currently leads a Sichuan Provincial International Science and Technology Innovation Cooperation project and collaborates with Hong Kong’s Child Assessment Service.

Li Sheng is a faculty member in the Department of Chinese and Bilingual Studies at the Hong Kong Polytechnic University. Previously, she held faculty positions at the University of Texas Austin and the University of Delaware. She directs the Language Learning and Bilingualism Lab, conducting research on language development mechanisms and improving quality of life for individuals with language disorders through developing diagnostic tools and intervention programs.

